# Climate drivers of malaria at its southern fringe in the Americas

**DOI:** 10.1101/674572

**Authors:** Karina Laneri, Brenno Cabella, Paulo Inácio Prado, Renato Mendes Coutinho, Roberto André Kraenkel

**Affiliations:** Grupo de Física Estadística e Interdisciplinaria, CONICET, Centro Atómico Bariloche, Bariloche, Río Negro, Argentina; Instituto de Física Teórica, Universidade Estadual Paulista - UNESP, São Paulo, SP, Brazil; LAGE do Departamento de Ecologia, Instituto de Biociências da Universidade de São Paulo, São Paulo, SP, Brazil; Centro de Matemática, Computação e Cognição (CMCC), Universidade Federal do ABC, Santo André, SP, Brazil

## Abstract

In this work we analyze potential environmental drivers of malaria cases in Northwestern Argentina. We inspect causal links between malaria and climatic variables by means of the convergent cross mapping technique, which provides a causality criterion from the theory of dynamic systems. Analysis is based on 12 years of weekly malaria *P. vivax* cases in Tartagal, Salta, Argentina—at the southern fringe of malaria incidence in the Americas—together with humidity and temperature time-series spanning the same period. Our results show that there are causal links between malaria cases and both maximum temperature, with a delay of five weeks, and minimum temperature, with delays of zero and twenty two weeks. Humidity is also a driver of malaria cases, with thirteen weeks delay between cause and effect. Furthermore we also determined the sign and strength of the effects. Temperature has always a positive non-linear effect on cases, with maximum temperature effects more pronounced above 25° C and minimum above 17° C, while effects of humidity are more intricate: maximum humidity above 85% has a negative effect, whereas minimum humidity has a positive effect on cases. These results might be signaling processes operating at short (below 5 weeks) and long (over 12 weeks) time delays, corresponding to effects related to parasite cycle and mosquito population dynamics respectively. The non-linearities found for the strength of the effect of temperature on malaria cases make warmer areas more prone to higher increases in the disease incidence. Moreover, our results indicate that an increase of extreme weather events could enhance the risks of malaria spreading and re-emergence beyond the current distribution. Both situations, warmer climate and increase of extreme events, will be remarkably increased by the end of the century in this hot spot of climate change.

## Introduction

Dynamics of malaria epidemics are strongly influenced by climate [1–3]. In particular, at the geographical fringes of its distribution, malaria dynamics are driven by environmental factors such as temperature, rainfall, and humidity [4], as well as epidemiological ones, such as immunity [5, 6]. In a high CO_2_ emission scenario, Northwestern Argentina, the studied region, will likely suffer a high temperature increase by the end of the XXI century, and the frequency of heat waves and intense rains will increase [7, 8]. All of these may impact parasite transmission rates [9], since they are modulated by mosquito vector abundance, and parasite survival and development, both strongly affected by climate. Malaria persistence in any given region requires a minimum set of environmental factors that allow both factors to be sustained [9, 10], and thus the quantification of the effects of climate is essential in order to predict, and help mitigate, the spread of the disease, especially under the current prospects of climate change [11, 12]. In particular, at the frontiers of malaria incidence, climatic factors fluctuate between values that curb malaria transmission and values that support it. This opens up the possibility of studying *in situ* the effects of climate on the disease dynamics.

In the Americas, the southern fringe of malaria distribution cuts through the north of Argentina, mainly in Salta province. About 99% of malaria cases there are caused by *Plasmodium vivax* parasite, although previously in the 20th century *P. falciparum* and *P. malariae* also circulated [13]. In particular *P. vivax* parasite has a liver dormant stage known as hypnozoite that causes relapses of the disease, i.e. secondary infections produced after a first infection [14]. Tropical *P. vivax* strains usually begin to relapse within a month after the initial infection, while hypnozoites of temperate strains usually have an incubation period of several months [14, 15]. For instance, in arid and semiarid regions of North-West India, relapses lead to two *P. vivax* seasonal peaks 8 months apart between each other [14].

The predominant malaria vector species in Salta province, northwestern Argentina, is the mosquito *Anopheles pseudopunctipennis* [16], although *An. argytarsis* and *An. strodei* are also found, together with other less abundant *Anophelines* [17]. Previous studies show that in the southern area of the subtropical mountainous rain forests in northwestern Argentina, the relative humidity is the major determinant of the abundance of *An. pseudopunctipennis*, *An. argytarsis*, *An. strodei* and *An. evansae* [17, 18]. Other studies claimed that maximum mean temperature and maximum mean humidity were the climatic variables that best explained the abundance fluctuation of *An. pseudopunctipennis* adults [19]. This species usually breeds in sun-exposed clean freshwater in association with floating plants and filamentous algae [20]. Its population density increases between December to April [21]. In some localities, three peaks of mosquito population were observed: the largest during the spring season (September-December), the second during autumn (March-May) and the smallest during the summer season (December-February) [19]. During winter, mosquito population drops to almost zero.

Local temperature is an important variable that determine mosquito survival but also parasite development inside mosquitoes. The Extrinsic Incubation Period (EIP), i.e. the time between successive infecting mosquito bites, depends strongly and non-linearly on temperature [9, 22]. Previous works performed in Kenya, Africa, report a range from 10 to 50 days for EIP of *P. falciparum* for temperatures in the range of 10°C − 37°C. Cooler temperatures give longer and more variable EIPs [22]. The EIP is often relatively long compared to the life expectancy of mosquitoes, which is around one month. The common assumption is that around 10% of the adult mosquito population will survive to successfully transmit the disease [23].

In Tartagal—the third most populated city in Salta Province, Northwestern Argentina, with 80,000 people—malaria cases occur typically from November to April (summer season), giving rise to an epidemic-like dynamics, characteristic of a disease at the limit of its geographical distribution. Most of the people infected with malaria work on agriculture in rural environments [16]. In fact, at the time that more workers are attracted to the region, serious flooding events increase the susceptibility [18, 24]; these are related to deforestation of the native forest near Tartagal, mainly to establish soy plantations. This trade-off between agricultural development and flooding that increases mosquito population, and therefore malaria risk, was addressed in other parts of the world [25] and might also impact malaria dynamics. In these low-incidence regions climate usually plays a key role in determining the seasonal pattern of malaria outbreaks. For instance, previous studies performed in El Oculto and Aguas Blancas, two localities situated approximately 100 km west from Tartagal, found an association between monthly malaria cases and maximum mean temperature and rainfall, or mean temperature and humidity, respectively [19].

Such correlations between climate and disease are often assessed in view of establishing how climate influences disease transmission. However, correlation has well-known drawbacks; most famously, “correlation does not imply causation”, because other factors, so-called confounding factors, may influence both variables. Furthermore, non-linearity may also obscure the relationship between the variables, to the point that even when there is a definite influence, correlation may not be observed, or can even change its sign—the “mirage correlation” problem [26]. In other words, “lack of correlation does not imply lack of causation”. Still, the goal is to determine whether climate drives disease dynamics; this led to new techniques aimed at detecting causal links from time series [27–30].

Convergent cross-mapping (CCM) is a method that tackles those issues considering the problem from a dynamical systems point of view. According to this view, two variables are causally linked if they are part of a coupled, possibly non-linear, dynamical system. This contrasts with the statistical criterion for association, either via direct correlation, or more complex statistical methods. Specifically, CCM assumes that dynamics are predominantly deterministic, and employs state space reconstruction, described in the methods section, to infer whether two variables are part of the same dynamical system, and therefore are causally connected [26]. Although it is based on classical theorems from deterministic dynamical systems, the method has been successfully tested on synthetic data [26, 31–33] and applied to a wide range of real-world problems [34–40].

Moreover, CCM is completely data-driven, that is, its results do not depend on choice of model equations or statistical models. Its results can also be used to quantify the direction and strength of the causal variables, e.g. environmental drivers, on the caused variable, such as number of cases, for each value of the causal variables [41]. This allows us to establish for which range of values the effect of a given variable is important.

We used CCM to analyze time series of 12 years of humidity, temperature, and malaria cases from Tartagal city. We established causal links between humidity and temperature, and malaria cases. We found that these effects lag the causal variable by periods compatible with some key biological components of malaria transmission.

## Materials and methods

Malaria *P. vivax* number of cases, provided by the Argentinean Ministry of Health, were recorded from January 2000 to December 2011 every epidemiological week. Symptoms of *P. vivax* are mild [16] but people that go to the hospital are treated with primaquine and chloroquine for 14 consecutive days [19]. Asymptomatics were not detected after specifically designed experiments that were carried out in order to fulfill the requirements to achieve the certificate of indigenous malaria-free country [42]. During the studied period, the National Program for Paludism Control (PNP) has developed activities of prevention and control consisting of active search for fever cases, and insecticide spraying every semester [16]. In that context, blood samples were taken from people with malaria symptoms, and blood smears were giemsa-stained for parasite detection. Positive samples were examined for identification of the *Plasmodium* species [19].

Tartagal city is located at the base of the Argentinean sub-Andean hills (22° 32’ S, 63° 49’ W; 450m above sea level) and holds 3/4 of the San Martín population department in Salta Province, Northwest of Argentina. The city is surrounded by subtropical native forests and crops such as beans, cotton, soybean, maize, grapefruit and tomato. The climate is subtropical, with an average annual temperature of about 23° C (maximum of 39° C in summer and average minimum of 9° C in winter). Annual cumulative precipitation is about 1100 mm, with a dry season from June to October with a monthly average rainfall of 30 mm, contrasting with the wet season from November to May with a monthly average rainfall reaching 160 mm [43]. It is possible to differentiate three seasons: one warm and dry, which corresponds to Spring (September-December), one warm and rainy, which corresponds to Summer (January-April), and one that is cold and humid which corresponds to Autumn and Winter (May-August) [17]. The city is crossed by the Tartagal River, and its urban area covers approximately 15 *km*^2^ [43]. Daily recorded climate data sets for Tartagal city for minimum and maximum relative humidity, rainfall, and minimum and maximum temperature were provided by the National Meteorological Service of Argentina. Rainfall data had too many missing values to perform the analysis, since the applied methodology requires short-time variability that cannot be recovered using seasonal means or data from other localities.

### Convergent Cross Mapping (CCM)

*Convergent cross-mapping* (CCM) is a method for detecting causality in nonlinear dynamic systems [26].

It is based on Takens’s theorem of *state-space reconstruction* finding that the attractor that governs a dynamical system can be retrieved through lagged coordinates (by multiples of a fixed period *τ*) of a single variable. The state-space reconstruction reveals a shape in an *E*-dimensional space, called *shadow manifold M*, which is an *embedding* of the dynamical system’s true attractor; in other words, it constructs an attractor topologically equivalent to the “original” one.

One of the corollaries of the generalized Takens’s theorem is the possibility of performing a cross-mapping between variables observed from the same system [44]. For example, consider two coupled variables, *x* and *y*, in a dynamical system, such that *x* affects *y*. The shadow manifold *M*_*y*_ based on *y* must map the corresponding (contemporaneous) value of the shadow manifold *M*_*x*_ based on *x*. CCM determines how well local neighborhoods on a given shadow manifold correspond to local neighborhoods on the other. From the reconstructed shadow manifolds, we can use *M*_*y*_ to predict states *x*_*p*_ in *M*_*x*_. The prediction skill is obtained by calculating the correlation coefficient *ρ* between predicted *x*_*p*_ and observed *x*_*0*_ states. Convergence means that the estimates from the cross-mapping improve as we increase the length of the time-series (*L*), because the larger the sample of the dynamics the better the shadow manifold portrays the true attractor. If there is causation, we expect to see convergence, since the correlation coefficient between predicted and observed increases as *L* increases.

Note that the causal variable leaves a signature on the affected one, but not *vice-versa*. This means that, if *x* causes *y* as above, *M*_*y*_ maps onto *M*_*x*_, but not the other way around, because *x* time-series contains no information about *y*.

In the dynamics of disease transmission, some particularities must be taken into account. For example, if temperature is a causal factor in malaria cases, then it is reasonable to expect that this relationship may not be instantaneous, since mosquito population dynamics and parasite cycle occur on the scale of several weeks. In this case, temperature may affect malaria cases with a delay, which we address by performing the CCM with increasing time lags between number of cases and temperature, and finding the delay with maximum prediction skill [31]. Of course, the same holds for other variables.

Another relevant aspect in this context is seasonality. Variables such as temperature and humidity are seasonal and so very predictable. Therefore, it is necessary to distinguish between the influence of the season and of the climatic variables themselves. This can be resolved by constructing a null hypothesis with time series surrogates [45]. For a given variable *x*(*t*) (e.g. temperature or humidity) we obtain a seasonal pattern *x*_*s*_(*t*) using smoothing splines, which works better than Fourier decomposition for non-sinusoidal waveforms. The residuals *x*_*r*_(*t*) are calculated by the difference between the seasonal cycle and the observed data *x*_*r*_(*t*) = *x*(*t*) − *x*_*s*_(*t*). These residuals are then shuffled and added back to the seasonal pattern, generating a surrogate series 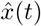 which has the same seasonality as the observed data but with shuffled residuals. Thus, if *x*(**t**) is really causally linked to another variable (e.g. *y*(*t*)) then *y*(*t*) should predict *x*(*t*) better than 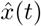. In other words, *y*(*t*) will be sensitive not only to the seasonal pattern but also to the anomalies (residuals) of the variable *x*. For each driver variable, we calculated the prediction skill *ρ*_*s*_ for 500 surrogates obtaining the probability distribution of *ρ*_*s*_. The value of prediction skill *ρ* for the original series was compared to the distribution of *ρ*_s_ to assess whether the result is statistically significant, using a significance level of *α* = 0.05.

Simple cross-mapping of attractors presents a few pitfalls; notably, high levels of noise and synchrony between the variables’ dynamics may lead to spurious conclusions [46]. We overcome the first issue by reestimating the number of malaria cases using a state space model, described in the next section. The problem with synchrony is dealt with in two parts: first we use a surrogate method, explained above, to make sure there is a causal link. Secondly, we analyze the time lag between the variables—a negative time lag is a reliable indication that the causal direction is correct [46–48].

### Quantification of interactions’ strength

Once we have established the causal variables, we investigated whether they have a negative or positive effect on the number of malaria cases using the *S map* multivariate approach [41]. It consists essentially of the same method of state space reconstruction, but including the causal variables into the state space axes. From that reconstruction, we recover a coefficient that is a proxy for the interaction strength between each driver and the target variable [41]. More specifically, consider that variable *y*(*t*) is affected by *E* different other variables: *x*_*i*_, *i* = 1, 2, … *E*. The state space at time *t* is given by: **x**(*t*) = *x*_1_(*t*), *x*_2_(*t*), … , *x*_*E*_(*t*), for each target time point *t*. The S-map method produces a local linear model *C* that predicts the value *y*(*t*) from the multivariate reconstructed state-space vector **x**(*t*) as follows:

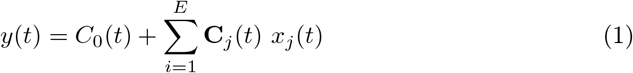

The model **C** contains the coefficients **C**_0_, **C**_1_ *…* **C**_*E*_ for each time point *t* and is obtained from the singular value decomposition solution according to the method presented in [41]. The coefficients **C**(*t*) are related to changes in the magnitude of causal factors over time and are used here to infer how malaria cases are affected by climate drivers.

### Expected number of malaria cases

Epidemiological time series usually show epidemic bursts separated by intervals with no cases recorded. Nevertheless the number of sequential zeros in a time series should be smaller than the dimension of the state-space reconstruction intended [45]. Moreover, an unknown number of cases are not detected, which introduces observational noise into the series. Observational and process noise can blur the causality links to be detected by CCM [44, 46].

To circumvent these problems we applied the CCM analyses to the expected number of recorded cases, which we estimated from the observed time series of number of cases, with minimal assumptions. To do that we fit to the time series a state-space model of count data dynamics in discrete time [49]. In this model the expected size of the population of infected people is the single state variable, and has a growth rate that can change at each time step. The number of infected people at each time step follow Poisson distributions, which have the expected number of cases as their only parameters. Finally, the expected number of recorded cases is a fixed proportion of the number of infected people that are detected. This model thus describes the number of recorded cases at each time step as a zero-inflated Poisson variable that evolves freely in time. Under this model, the observed time series of recorded cases is a realization of a stochastic process, from which we estimate the expected trajectory in time. Therefore, the time series of expected number of cases estimated by this model sorts out observation noise caused by detection failures and also averages out process noise.

We used a Bayesian fit of the model to the data with a Markov Chain Monte Carlo (MCMC) in JAGS [50] through the *rjags* [51] and *R2jags* [52] R packages. We ran four MCMC chains of 3.6 10^6^ interactions each, with a burn-in period of 1.51 10^6^ interactions, and thinning interval of 1500 interactions, which thus returned a sample of 4, 000 values from the posterior distributions of the expected number of cases at each time step. Chain convergence was checked by the 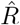 statistic, which had a maximum of Effective sample size of posteriors were 560. Codes for the fitting and binary files with the complete output are available at https://github.com/mathbio/malariaCCM.

### Analyses performed

We applied the CCM analysis to the time series of expected number of malaria cases being caused by time series of maximum and minimum weekly averaged temperatures, maximum and minimum temperatures’ weekly standard deviation, maximum and minimum weekly temperature amplitudes, and maximum and minimum weekly averaged relative humidities. Each variable was tested for lagged effects up to 30 weeks, since we intended to capture the causality relationships in a single epidemic and mosquito cycle. As previously shown [31], several secondary time-lags may also be causally related as a by-product of the main lag. In this case, we chose the significant time-lag with maximum prediction skill *ρ*, and used it to determine the signal and strength of the effects.

## Results

Within the studied interval (02 Jan 2000 – 11 Dec 2011), a total of 266 cases of malaria caused by *P. vivax* were recorded in Tartagal, with an average of 0.43 case per week and a maximum of 16 cases in a single week. Malaria cases in Tartagal are typically observed from November to April (summer season), while during winter (July to September) they drop to zero over several months (Fig. 1).

**Figure 1.**
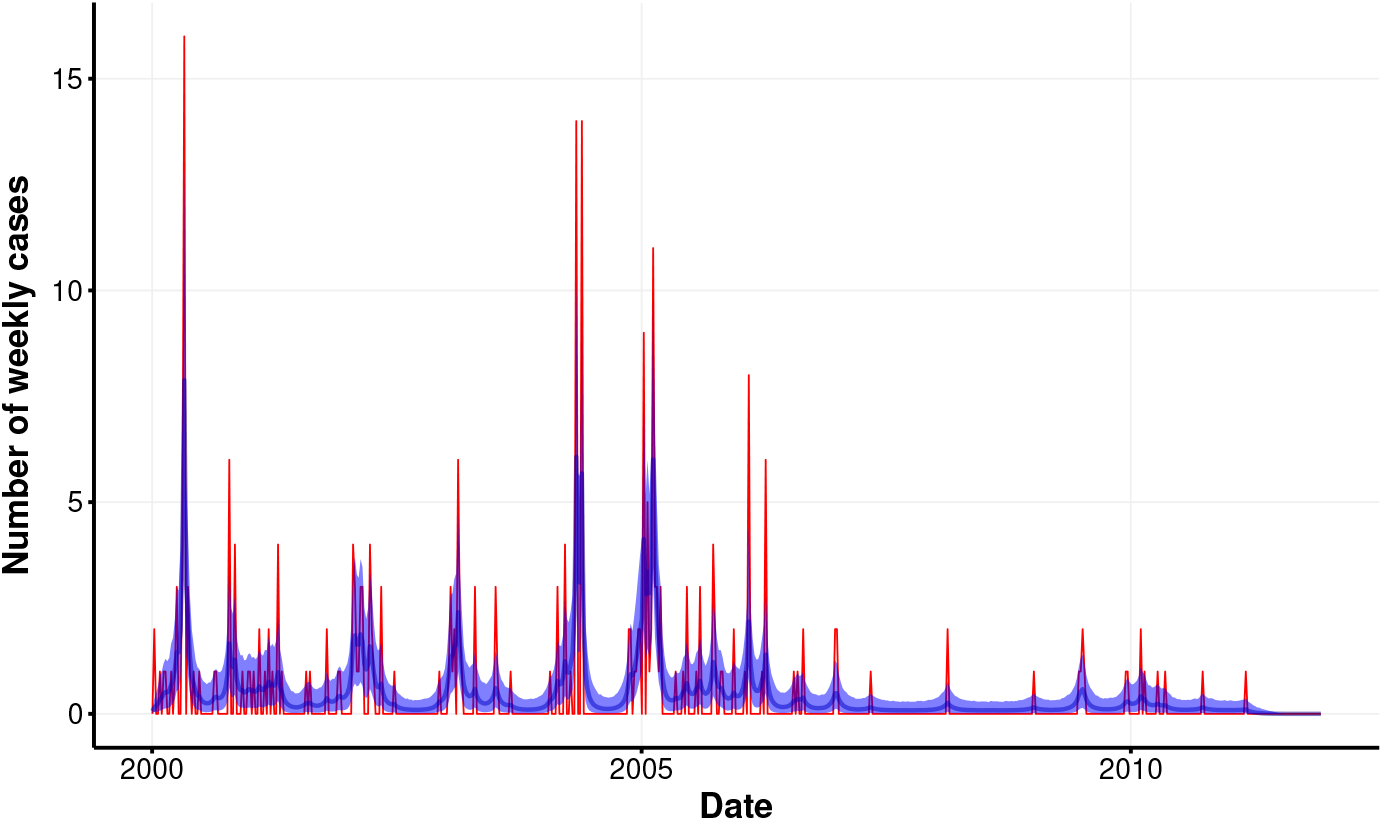
Weekly malaria *P. vivax* cases for Tartagal city (red) and estimated mean weekly number of cases with its 95% credibility interval (blue).

The estimated mean weekly number of cases were compatible with the observed counts, ranging from 4.5 × 10^−5^ to 7.88 (mean of posterior distribution), following the same temporal trend of the observed number of cases (Fig. 1). The number of observed cases were in general larger than the estimated averages, which is expected for count variates with low mean values.

### Seasonality

Temperature in Tartagal ranged from 0° C to 39.8° C with a mean of minimum temperatures of 16.1° C and mean of maximum temperatures of 27.8° C, and humidities ranged from 18.1% to 98.7%. All climate variables showed a clear seasonal trend, that express the alternation of a rainy, warm and wet summer (December – February) to a colder and not so wet winter (June – August, Fig. 2).

**Figure 2.**
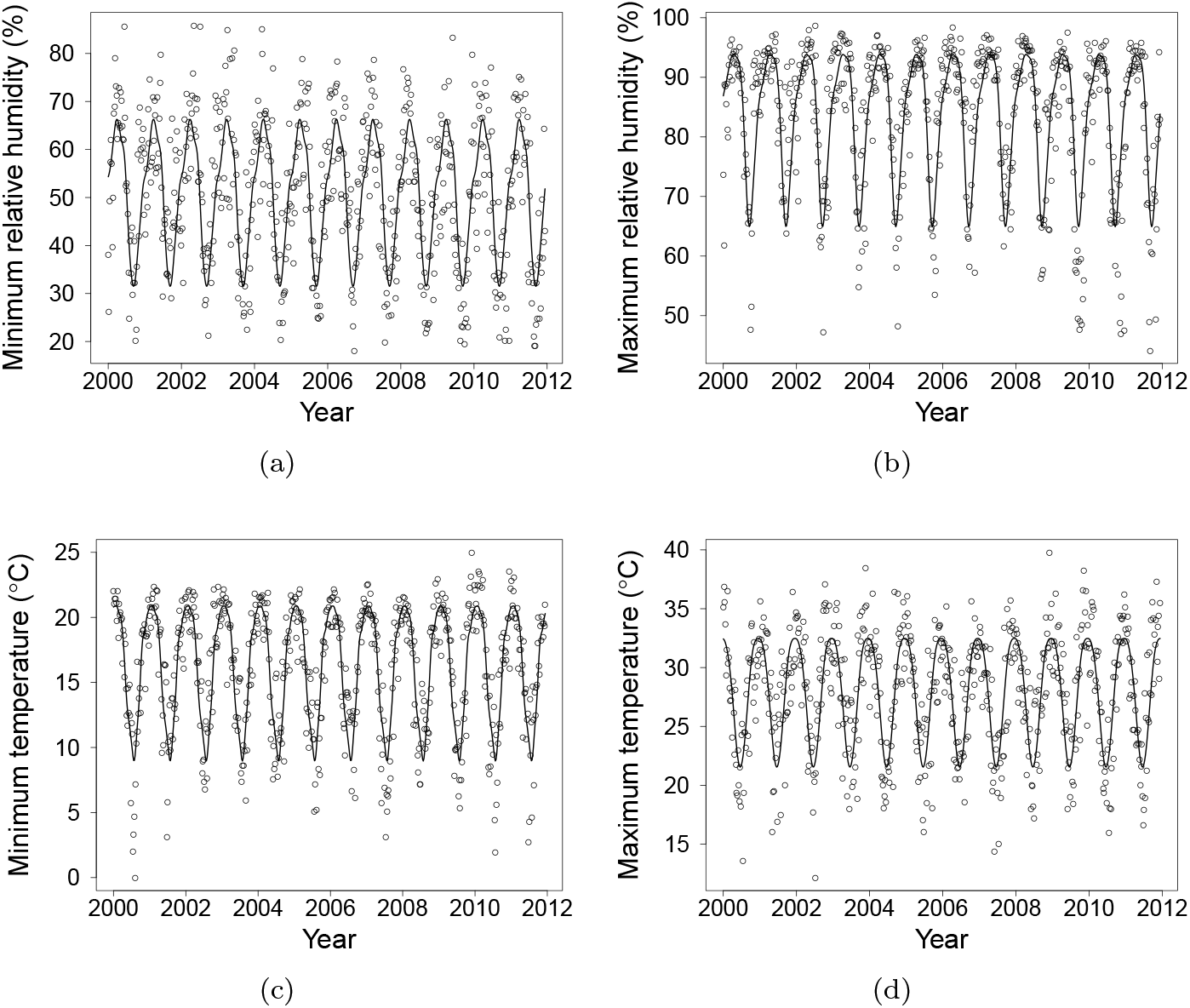
Weekly values of climate variables (points) and their seasonal trends, fitted by spline regressions.

### Causality tests

We found causal relationships between four climate variables and malaria cases in Tartagal. Maximum temperature was causally linked to expected number of malaria cases 5 weeks later, while minimum temperature, around both 0 and 22 weeks of time-lag, was also important. Maximum and minimum humidity showed causal links to expected number of cases as well, with a lag of *ca.* 13 weeks (Fig.3). In all cases, the prediction skill (*ρ*) was about 0.5, and there were several time-lags at which the causal link was significant. As justified in the methods section, we picked the time-lags with maximum *ρ* to interpret the result and perform the following analyses. The CCM analysis showed non statistically significant results for both maximum and minimum temperatures’ standard deviation, and maximum and minimum temperature amplitudes, for the whole range of time-lags from 0 to 30 weeks.

**Figure 3.**
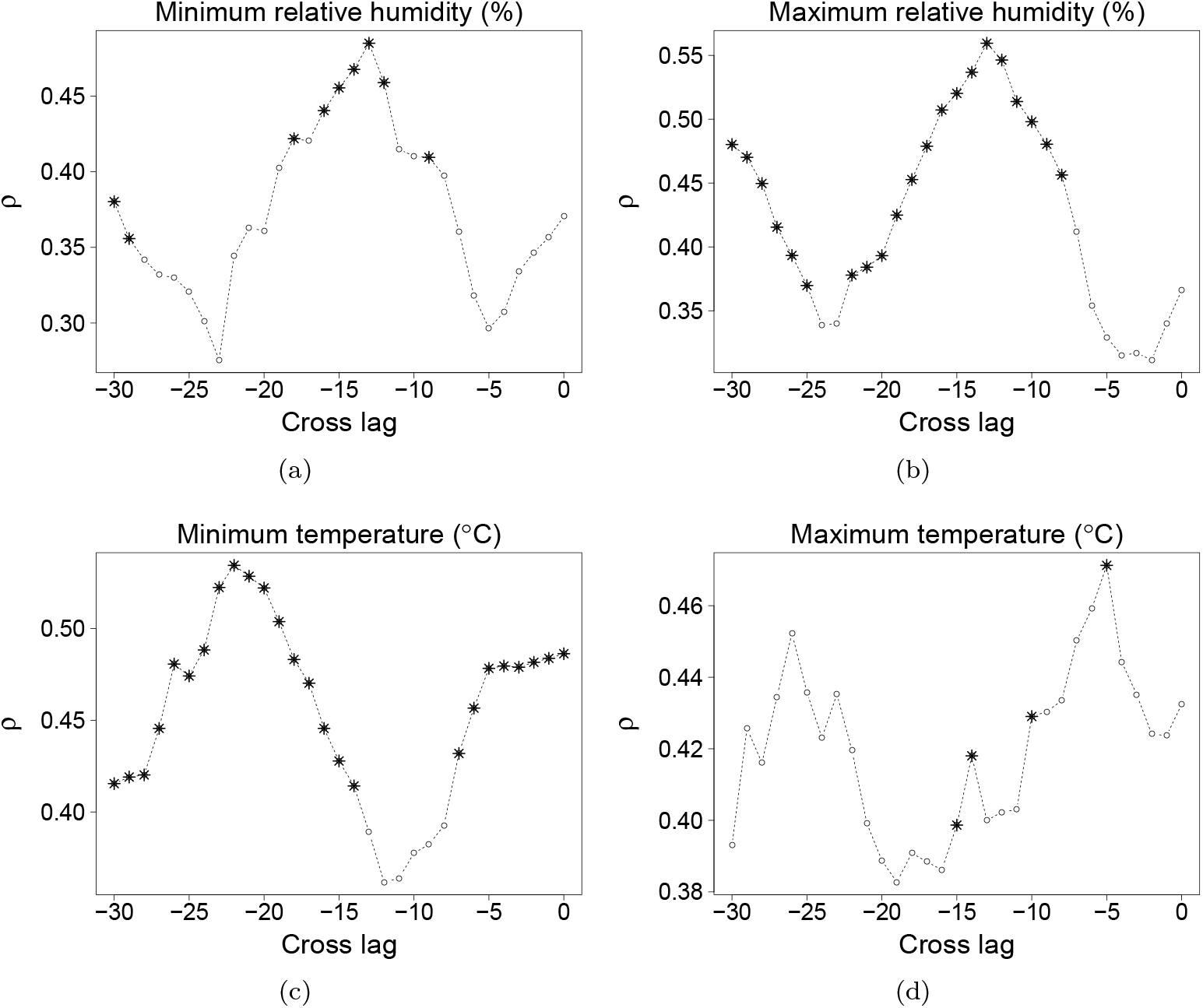
Significant causal relationships as a function of time-lag via CCM. Plots show in *x*-axis the time-lags tested for each climate variable and in *y*-axis the prediction skill *ρ* of the observed mean number of malaria cases. Asterisks represent time-lag values that provided a larger *ρ* than expected by a common seasonal trend, considered significant with *α* = 0.05.

We note that correlation was weak between the climatic variables and number of malaria cases: the correlation coefficients are, at most, of 0.3 for all variables and lags studied, systematically below the values of prediction skill *ρ* found. This means that, in our study, correlation would not be enough to detect the patterns we found.

### Interactions’ strength and direction

Here we look at each causal variable found above and analyze the effect it has on the number of malaria cases (Fig. 4). As expected for non-linear dynamics, both the strength and direction of the causalities that we detected changed markedly along the time series, showing that the causal effects can emerge, wane, and even reverse as this complex dynamics unfolds. Minimum humidity lagged by 13 weeks has a positive effect on number of malaria cases almost all over the time series, but the strength of the interaction decreases with increasing values of minimum humidity (Fig. 4 a), that is, an increase in this variable has a stronger positive effect when minimum humidity is low. In contrast, maximum humidity effect (also lagged by 13 weeks) has generally a negative effect, which becomes more pronounced for values above 85% relative humidity.

**Figure 4.**
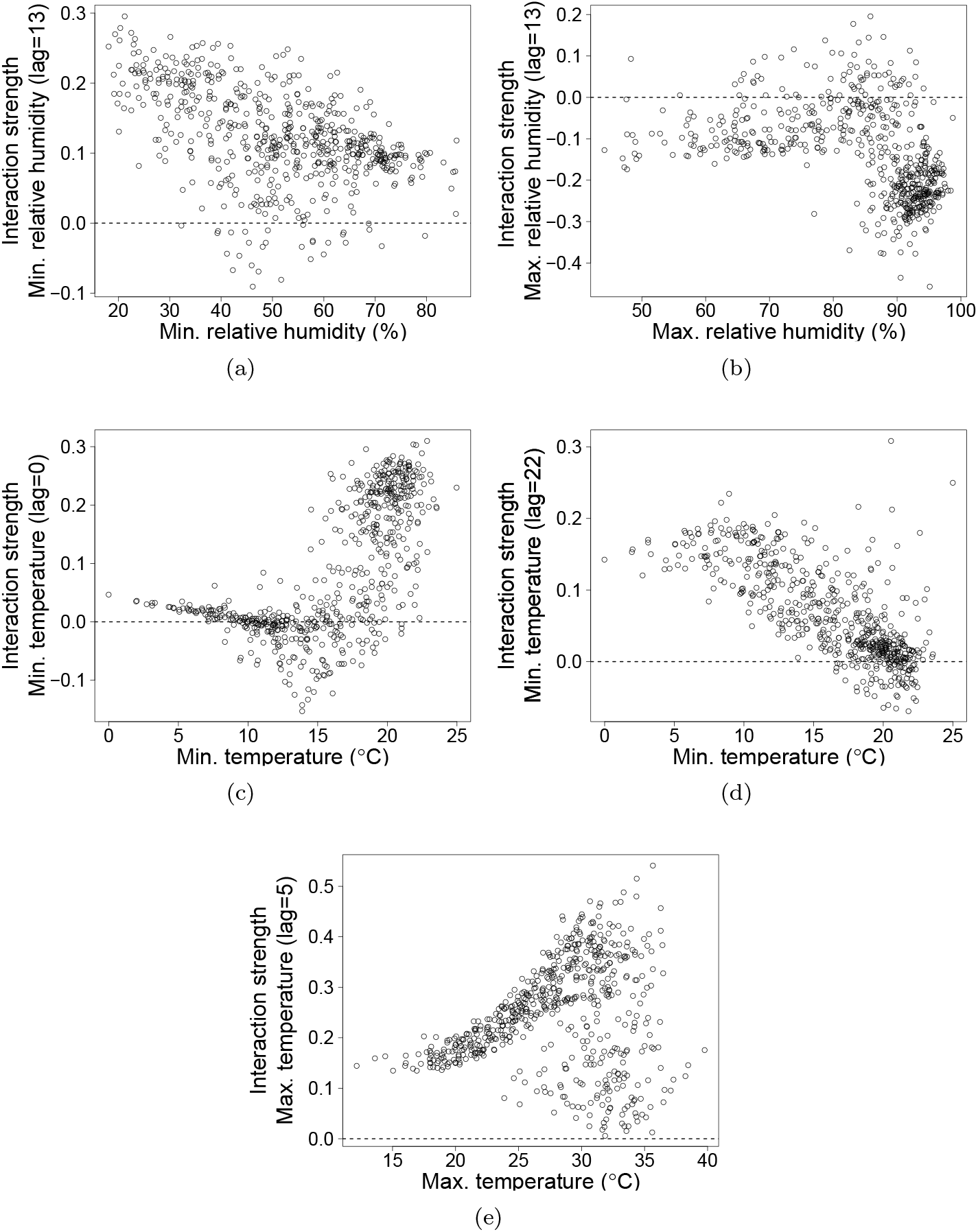
Interaction strengths for each causal variable as a function of the cause itself. (a) Minimum relative humidity with time-lag of 13 weeks has a positive effect on the number of malaria cases in most of the points of the time series, though this causality weakens as humidity increases. (b) Maximum relative humidity with time-lag of 13 weeks had in general a negative effect on cases that is stronger above 85% of humidity. (c) Minimum temperature with no lag tends to affect positively the number of cases, especially above 15° C. (d) Minimum temperature lagged by 22 weeks had the same pattern as minimum relative humidity in (a). (e) Maximum temperature lagged by 5 weeks also had positive effect on cases in most of the time series, with increasing causal strength as maximum temperature raises.

These two results show that cases of malaria are bounded by relative humidity lagged by 13 weeks, which has stronger effects close to the extremes of humidity recorded for the region (around 20-30% minimum humidity and above 85% maximum humidity, Fig. 2 a-b).

In general a rise in temperature causes an increase in the number of cases, although the strength of this effect is contingent on current and lagged temperatures. The increase of minimum temperature at lag zero (in the same week as registered cases) had mainly a positive effect on the number of malaria cases when minimum temperature was above 17° (Fig. 4), and such effect strength increases abruptly when minimum temperature increases. The effect of the minimum temperature lagged by 22 weeks, on the other hand, gets weaker with increasing temperature, even though its effect was also usually positive. Finally, maximum temperature lagged by 5 weeks always had a positive effect on number of cases, an effect that was stronger and much more variable above 25° C.

The thresholds for the interaction strength were checked by comparing the mean effect below and above the assumed threshold using a Mann-Whitney (Wilcoxon rank sum) test. In all cases, the test rejected the null hypothesis (*p* < 2.2*e* − 16 for all thresholds).

## Discussion

We used a causality criterion from dynamic systems theory to identify causal relationships between climatic factors and number of malaria cases in Northwestern Argentina. With this approach we were able not only to quantify causal effects (summarized in Table 1) but also to highlight key features of the disease dynamics without the need of an explicit model.

**Table 1.** Summary of significant causal variables found for malaria cases, the direction of their effects, and the behavior of the relationship with different values of the variable.

| variable | lag (weeks) | sign | range of values |
| --- | --- | --- | --- |
| Min. relative humidity | 13 | positive | decreases with humidity |
| Max. relative humidity | 13 | negative | pronounced above 85% |
| Min. temperature | 0 | positive | pronounced above 17° C, increases with $T$ |
| Min. temperature | 22 | positive | decreases with temperature |
| Max. temperature | 5 | positive | more pronounced and variable above 25° C |

In a nutshell, we show that causal links to climatic variables occur at different time lags with respect to number of cases, and can be interpreted as pertaining to two factors for the transmission of the disease. distinct scales: a longer one relevant for mosquito population dynamics, and a shorter one compatible with pathogen cycle. These processes can bring on high vector abundance and conditions favoring parasite development, which are *sine qua non*

Number of cases was causally linked to minimum and maximum temperatures lagged by 0 and 5 weeks respectively, with increasing temperature leading to higher number of cases. These short time lags are probably linked to the Extrinsic Incubation Period (EIP) of the malaria parasite inside the vector, which strongly depends on temperature [22, 53, 54]. Malaria symptoms, and therefore cases, appear approximately one week after EIP completion: some works report a range from 10 to 50 days for EIP [9, 22], taking longer for lower temperatures. Therefore, lower temperatures can quickly interrupt the parasite cycle, as it becomes longer than mosquito lifespan, in agreement with the immediate (zero time lag) response of minimum temperature, which becomes more sensitive around 17° C. This concurs with recent estimates [9] for *P. falciparum*, which is better studied than *P. vivax*, but has a similar dependence on temperature [54]. On the other hand, increasing temperatures will reduce EIP and increase the rate of transmission, leading to higher numbers of cases 10 to 50 days later, which is compatible with the lag of 5 weeks found for maximum temperature.

At larger time lags, we found that minimum and maximum humidities, with lag of 13 weeks, and minimum temperature, with lag of 22 weeks, were causally connected to number of malaria cases. The effects were positive for minimum humidity and negative for maximum humidity, indicating that intermediate values of humidity lead to a higher number of malaria cases. The lagged effect of minimum temperature was positive and especially strong for low temperatures, below 15° C. In other words, at these temperatures but 22 weeks before reported cases, an increase in minimum temperature has a strong positive effect on the number of cases. These long time lags could influence the dynamics of mosquito population, which are a key component of the malaria transmission. Data obtained from laboratory experiments with controlled temperatures and humidities show that low humidities are always detrimental to adult mosquito survival [55]. On the other hand, these same data show survival was maximal at 100% humidity for temperatures up to 20° C, while for temperatures above 25° C (inclusive) it was slightly higher for 80% humidity than 100% (see Fig. 6.1 in [55]). It was also found that low temperature curbs population growth by halting larval reproduction [56]. Moreover, an observational study in Aguas Blancas and El Oculto, in the same region of Argentina where the analyzed data was collected, showed that mosquito population usually peaks around October, before the peak of malaria cases [19]. This means that vector abundance does not immediately incur a high number of cases, hence the actual factor in the transmission dynamics is not a large mosquito population size *per se*, but the maintenance of population, and possibly parasite density, at certain levels—thereby, during the period favorable to the malaria parasite there will be a mosquito and human population suitable to maintain the transmission cycle. Accordingly, the time lag between climatic factors affecting malaria parasite, mosquito population dynamics and number of cases can indeed be very long.

Along this line, the mentioned studies [19] show that *Anopheles* mosquito population drops almost to zero during winter, except in the last year of the study, when mosquito population remained low. Therefore, it is expected that transmission goes down abruptly in the winter season. This fact, together with the long extrinsic incubation period during winter, makes transmission almost halt from June to August. On the other hand, some of the cases reported during this period might be relapses. Since these are not well documented in Argentina, we performed an exploratory autocorrelation assessment, using the same methods as [14], detailed in the Appendix. It provides some evidence that relapse cases are widely distributed between 4 to 10 months for the time series we analyzed (Fig.A1). This means that relapse cases should not affect the interpretation of the time lags above.

The importance of weather and climate in malaria outbreaks at the edge of its geographical distribution has been supported by the good fit and the predictive power of epidemiological models that incorporate climate variables, e.g. [3, 4, 57]. This statistical modeling approach gauges the relevance of a candidate causal variable by the likelihood of the models that include some effect of the variable [58, 59]. Therefore, the criterion to infer a causal link must assume a function to describe the cause-and-effect relationship. On the other hand, CCM does not rely on an explicit model, but on reconstructions of the attractor as data-driven descriptions of the dynamics from which the observed time series come. In this approach causality implies that the affected variable’s attractor can be mapped onto the causal variable’s attractor. This cross-mapping does not depend of the specific way one variable affects another, and is not affected by changes over time of the interaction between variables, which commonly occur in non-linear dynamics.

In other words, CCM spots causal links even if the effects of the causal variables are contingent on other (possibly unknown) variables through unknown functions. Statistical models can also take into account such contingencies by making them explicit in interaction terms of linear models or with nonlinear functions. Nevertheless, even a modest set of candidate causal variables ensues a huge number of alternative combinations of variables, lags and functions to be evaluated. Sorting out causal links from so many alternatives can turn into a “data-dredging expedition”, and can also end up with over-parametrized models or with an uninformative set of alternative models [58]. For the sake of elucidating causes of a phenomenon—as was our goal—CCM avoids these pitfalls because it is model-free. Therefore, besides its value in providing an alternative criterion to detect causal links, CCM can also be used to select variables to be included in predictive statistical or dynamical models, as well as to choose the best functions that describe causal links in these models.

In summary, we have applied CCM to detect and characterize causal links between environmental factors and number of malaria cases using epidemiological and climatic data. The use of observational data (time series of recorded cases and climate variables) brings together the dynamics of both vector and parasite transmission cycles, yielding a more comprehensive picture of the epidemiology of malaria, which is difficult to infer from laboratory studies alone. In this context, the study of malaria cases in the southernmost region of malaria incidence in the Americas allowed us to infer the climatic factors, and thereby the processes, that prevent the re-emergence of the disease beyond the present region.

Additionally, our analyses highlighted some less appreciated consequences of climate warming and increasing climate fluctuations that future models can take into account. We have shown that increased maximum temperature not only increases the number of cases of malaria, but also that this temperature effect accelerates as temperature rises. This non-linear effect makes warmer areas prone to more severe increases in disease spread with increasing temperatures, such as the tropical zones where malaria is or has been endemic. Moreover, the spread of disease would be facilitated by any climatic change that increases the probability of weeks with minimum temperature above its average (16°C in Tartagal). Also, anomalous wet weeks during the dry season or dry weeks in the rainy season cause an increase in the number of cases in the following weeks. Therefore, an increase of extreme weather events also enhances the risks of malaria spreading and re-emergence beyond the current geographical distribution. According to climate model predictions [8], Northwestern Argentina is among the places that will suffer the highest increase of temperature by the end of the century. The causal links proposed in this paper can contribute to modeling future scenarios of malaria and other vector-borne diseases re-emergence in this hot spot of climate change.

## Acknowledgments

We thank two anonymous reviewers for their careful reading and helpful comments on the manuscript. We thank the Ministry of Health of Argentina and the National Meteorological Service of Argentina for providing the datasets. K.L. is member of CONICET, Argentina. We thank the ICTP-South American Institute for Fundamental Research (ICTP-SAIFR) for support, via the FAPESP grant #2016/01343-7. B.C. thanks Capes for support through a PNPD fellowship. R.A.K. and P.I.P. thank CNPq (Brazil) for partial support. R.M.C. thanks FAPESP for partial support through a fellowship (grant #2014/23497-0).

## Appendix

Relapse cases of *P. vivax* in Argentina have not been well documented. For other subtropical regions with low transmission, the period between relapses can be very long, from 8 months to years [14, 42, 60]. Also in low transmission settings the number of relapses due to a primary infection tend to be small, from 1 to 4 relapses [60]. In Argentina there are no data on relapse times—patients that are treated after the first infection are not followed up to record when they experience relapses. Parasitemia levels have not been measured either. Even if *P. vivax* parasitemia levels were much lower than in the case of *P. falciparum*, parasitemia increases with relapses but immunity also increases.

As a preliminary exploration, we estimated the timing of relapse cases by assessing the autocorrelation of the number of malaria cases time series [14]. Even though there are few cases in Argentina and a high associated measurement error, we obtained a slight autocorrelation increase from 4 to 10 months, which is in agreement with previously reported results of *P. vivax* in India [14]. For a more accurate measure of relapse times, future studies would be needed to follow up cases.

**Figure A1.**
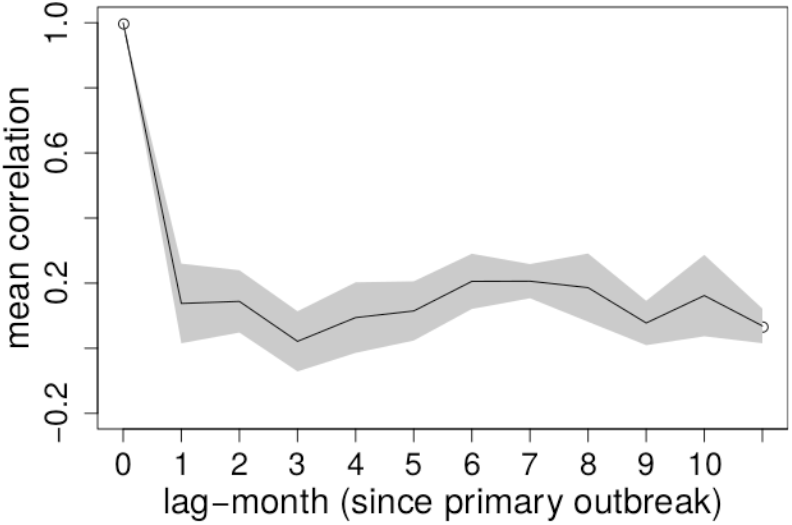
Averaged auto-correlation of *P. vivax* malaria cases (solid line) and associated error (gray) for each of the transmission months (November-April) with malaria cases in the subsequent 12 months. It slightly increases between 4 to 10 months from the primary outbreak, associated with relapses period. The average was done over all years.

